# Energy dynamics for systemic configurations of virus-host co-evolution

**DOI:** 10.1101/2020.05.13.092866

**Authors:** Alessandra Romano, Marco Casazza, Francesco Gonella

**Affiliations:** Scuola Superiore di Catania, Università degli Studi di Catania, Via Valdisavoia, 9, 95123, Catania, Italy; Dipartimento di Chirurgia e Specialità medico chirurgiche, CHIRMED, Università degli Studi di Catania, Via Santa Sofia, 78, 95124, Catania, Italy; Division of Hematology, AOU “Policlinico - Vittorio Emanuele”, Via Santa Sofia, 76, 95123, Catania, Italy; Department of Molecular Sciences and Nanosystems, Ca’ Foscari University, Via Torino 155, 30170, Venezia Mestre, Italy; Research Institute for Complexity, Ca’ Foscari University, Dorsoduro 3246, 30123, Venice, Italy

**Keywords:** Systems Thinking, simulator, positive-strand RNA virus, COVID-19, target therapy, systemic drugs

## Abstract

Virus cause multiple outbreaks, for which comprehensive tailored therapeutic strategies are still missing. Virus and host cell dynamics are strictly connected, and convey in virion assembly to ensure virus spread in the body. Study of the systemic behavior of virus-host interaction at the single-cell level is a scientific challenge, considering the difficulties of using experimental approaches and the limited knowledge of the behavior of emerging novel virus as a collectivity. This work focuses on positive-sense, single-stranded RNA viruses, like human coronaviruses, in their virus-individual host interaction, studying the changes induced in the host cell bioenergetics. A systems-thinking representation, based on stock-flow diagramming of virus-host interaction at the cellular level, is used here for the first time to simulate the system energy dynamics. We found that reducing the energy flow which fuels virion assembly is the most affordable strategy to limit the virus spread, but its efficacy is mitigated by the contemporary inhibition of other flows relevant for the system.

**Summary:** Positive-single-strand ribonucleic acid ((+)ssRNA) viruses can cause multiple outbreaks, for which comprehensive tailored therapeutic strategies are still missing. Virus and host cell dynamics are strictly connected, generating a complex dynamics that conveys in virion assembly to ensure virus spread in the body.

This work focuses on (+)ssRNA viruses in their virus-individual host interaction, studying the changes induced in the host cell bioenergetics. A systems-thinking representation, based on stock-flow diagramming of virus-host interaction at the cellular level, is used here for the first time to simulate the energy dynamics of the system.

By means of a computational simulator based on the systemic diagramming, we identifid host protein recycling and folded-protein synthesis as possible new leverage points. These also address different strategies depending on time setting of the therapeutic procedures. Reducing the energy flow which fuels virion assembly is addressed as the most affordable strategy to limit the virus spread, but its efficacy is mitigated by the contemporary inhibition of other flows relevant for the system. Counterintuitively, targeting RNA replication or virion budding does not give rise to relevant systemic effects, and can possibly contribute to further virus spread. The tested combinations of multiple systemic targets are less efficient in minimizing the stock of virions than targeting only the virion assembly process, due to the systemic configuration and its evolution overtime. Viral load and early addressing (in the first two days from infection) of leverage points are the most effective strategies on stock dynamics to minimize virion assembly and preserve host-cell bioenergetics.

As a whole, our work points out the need for a systemic approach to design effective therapeutic strategies that should take in account the dynamic evolution of the system.

## Article

Positive single-stranded RNA ((+)ssRNA) viruses, including picornaviruses, flaviviruses, togaviridae and human coronaviruses (h-CoVs) ^1-4^ cause multiple outbreaks, for which tailored antiviral strategies are still missing ^5-7^. (+)ssRNA viruses package their genomes as messenger sense, single stranded RNA and replicate those genomes solely through RNA intermediates in the cytosol of the host cells ^8^. RNA-dependent RNA polymerases lack co- and post-replicative fidelity-enhancing pathways, and final RNA genome copies incorporate mutations at a much higher rate than that observed for DNA genomes ^9,10^ providing the viral *quasispecies* a higher probability to evolve and adapt to new environments and challenges during infection ^11,12^. The diversity is essential for both viral fitness and pathogenesis, due to the complex relationship between virus replication, host cells and immune system, since almost +RNA viruses can delay antiviral innate immune response ^13^ in multiple ways ^14-20^. Host immunogenetic factors can be sensitive to a variation in the viral load, conveying to a defective response of the innate immunity that could explain the variable clinical course of infection ^20,21^.

Among (+)ssRNA viruses, h-CoVs have unusually large genomes (∼30 kb) associated with low mutation rates ^20,22^. Three novel h-CoVs are etiological agents of serious viral interstitial pneumonia ^23^. They include the severe acute respiratory syndrome, first described in Guandong province in China in 2002 ^24^, the Middle East respiratory syndrome (caused by MERS-CoV), emerged in June 2012 in Saudi Arabia and Qatar ^25^, and the SARS-CoV-2, emerged in China in January 2020, cause of the largest outbreak in the modern history ^23^.

Recent studies confirmed the complexity of viral dynamics, whose fitness is improved by the complex interactions with the host proteins ^26,27^ as previously described in modelling the virus-host interactions at sub-cell and cell levels ^6,28-31^. However, models that address a specific aspect of the virus-host interaction do not capture the wide range of intertwined spatial and temporal (hours to days) dynamic scales ^32^, that are related to the interaction of different concurrent hierarchical levels. All these factors address the need for a description able to connect all the levels at which the infection proceeds within a single cell, from the recruitment of RNA-synthesis factory to the final host cell damage. Several drugs, such as chloroquine ^33,34^, hydroxychloroquine, ribavirin, lopinavir-ritonavir ^35^, were used in patients with SARS ^33-35^ or MERS ^7^ and are currently under investigation, alone or in combination, to control the COVID-19 infection ^36-47^. However, there is an urgent need of a new class of compounds, other than nucleotides mimetics, due their low-fidelity viral RNA polymerase ^48^ to combine with host factor targeting agents, recently identified to lead to a therapeutic regimen to treat COVID-19 ^6^.

We show that, by approaching the viral infection as a dynamic systemic problem ^49,50^, it is possible to identify effective systemic leverage points to minimize the release of virions and virus spread in the body. We address this knowledge gap using a systems-thinking based simulator to build up a systemic description of the viral action, able to follow changes in the energy dynamics of the interaction between a (+)ssRNA virus and the host cell and therefore addressing effective intervention strategies.

### Systems-thinking diagram of the virus-host interaction

Starting from the knowledge of relevant processes in (+ss)RNA virus replication, transcription, translation, virions budding and shedding and their energy costs (reported in **Supplementary Methods Table 1**), we built up a systems-thinking (ST) based energy diagram of the virus-host interaction. **Figure 1** shows the stock-flow diagram for the system at issue, where each stock was quantified in terms of embedded energy of the corresponding variable. Symbols are borrowed from the energy language ^51,52^: shields indicate the stocks, big solid arrows the processes and line arrows the flows, whereas dashed lines indicate the controls exerted by the stocks on the processes. All stocks, flows and processes are expressed in terms of the energy embedded, transmitted and used during the infection. We used ATP-equivalents (ATP-eq) as energy unit ^53^ referred to cellular costs, by using the number of ATP (or GTP and other ATP-equivalents) hydrolysis events as a proxy for energetic cost ^53,54^. The dynamic determination of flows was based on the knowledge of characteristic time-scales of well-established biological processes (for more details see **Supplementary Methods**). The output flows *J*_*4*_ and *J*_*5*_ were set to be effective only if the value of the respective stocks *Q*_*3*_ and *Q*_*5*_ is higher than a threshold, as represented by the two switch symbols in the diagram.

**Figure 1.**
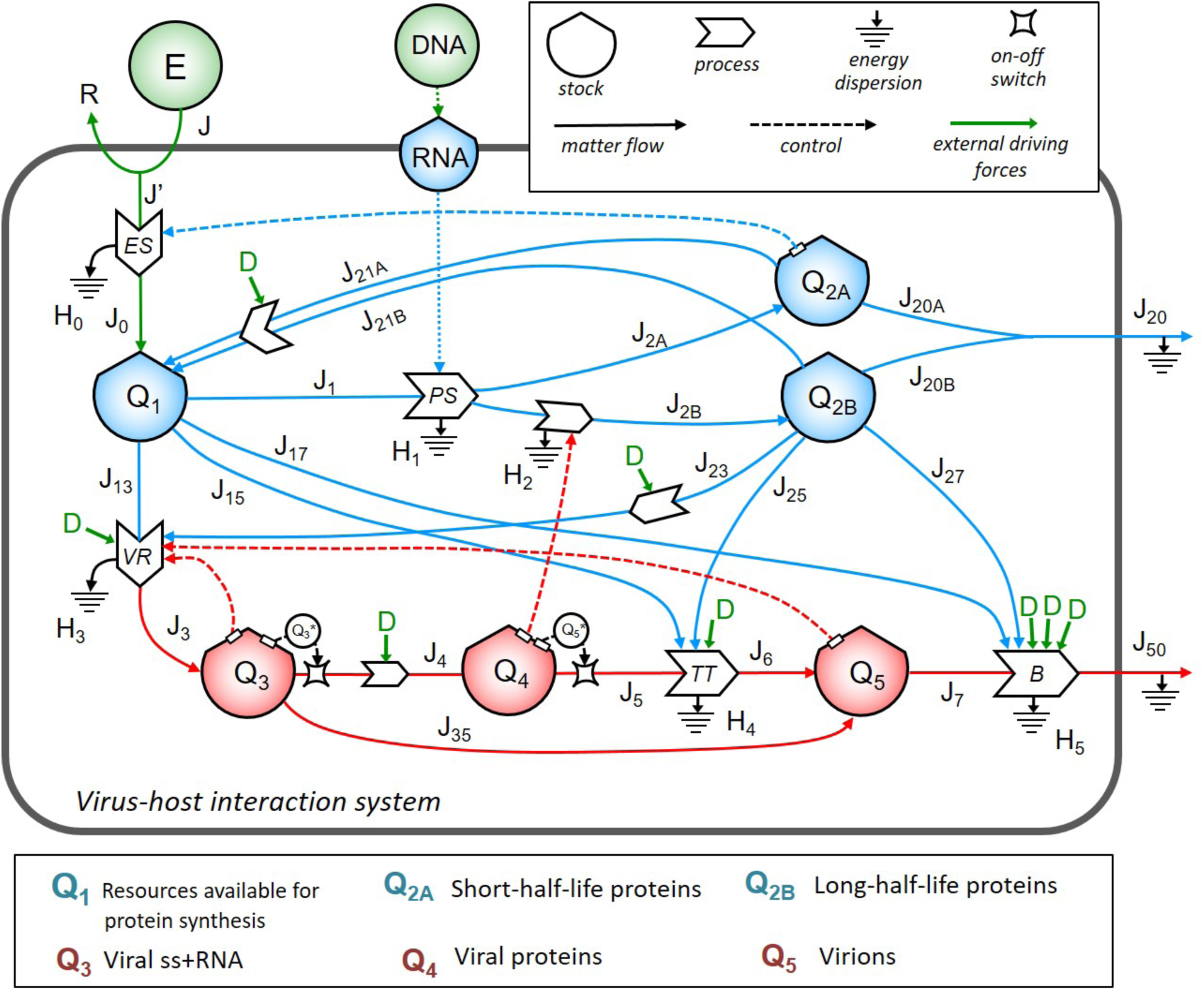
The energy systemic diagram of a cell infected by ss+RNA virus. Stock-flow diagram of the virus-host interaction system. In the upper right box are the meaning of symbols. The color code is: blue for host cell energy stocks and relative inflows and outflows; red for virus energy stocks and relative inflows and outflows; green for external energy inputs and external driving forces *F* corresponding to different therapic strategies. The lower box lists the biological contents of the stocks, all expressed in terms of energy (ATP-equivalent units).

The stock *Q*_*1*_ represents the embedded energy of resources addressed to protein synthesis in the host cell. The dynamics of allocation for protein synthesis depends on the cell bioenergetics, e.g., the number of mitochondria, OX-PHOS activity levels, and the cell cycle phase ^55-57^ (for more details, see **Supplementary Methods**). In the absence of virus, energy flows from *Q*_*1*_ (flows *J*_*1*_, *J*_*2A*_ and *J*_*2B*_) to produce short-half-life proteins (stock *Q*_*2A*_) and long-half-life proteins (stock *Q*_*2B*_), whose synthesis, degradation and secretion follow dynamic stationary conditions. In particular, typical of the specialized cells such as those of the pulmonary epithelium, there is a flow of proteins destined for degradation and recycling through the recruitment of autophagic receptors or organelles or proteasomes (identified in the diagram by the flows *J*_*21A*_ and *J*_*21B*_, **Supplementary Methods Table 1**). The outflow of folded, fully functional proteins addressed to secretion or surface exposure is represented by flows *J*_*20A*_ and *J*_*20B*_, grouped into the flow *J*_*20*_. At the time of infection, the presence of viral load in the system, expressed by the stocks *Q*_*3*_ (viral RNA content to be used for viral transcription and translation), *Q*_*4*_ (translated viral proteins content) and *Q*_*5*_ (full assembled virions to shed virus outside) diverts resources directly from *Q*_*1*_ (through flows *J*_*13*_, *J*_*15*_ and *J*_*17*_) and *Q*_*2B*_ (through flows *J*_*23*_, *J*_*25*_ and *J*_*27*_). Virions shedding is represented by the flows *J*_*7*_ and *J*_*50*_ through the contribution of the host-flows *J*_*17*_ and *J*_*27*_ (see **Figure 1**).

### Dynamics of a cell infected by a +ssRNA virus

From the energy ST diagram, we derived the equations to set up a computer simulator of the system (**Supplementary Methods Table 1**). Each flow was represented by an equation to explicit the dependence of the processes on the various stocks. To this end, we applied the standardized workflow of systemic modeling (**Extended Data Figure 1**) ^51^, typically used to model the dynamics of resources within a complex system ^58^. Calibration values were determined for the set of *Q*_*i*_ initial conditions and the set of phenomenological coefficients *k*, derived from established characteristic time-scales for each process in the diagram (further details and references are available in **Supplementary Methods**).

First, we investigated the system dynamics under different initial conditions, exploring the possible role of different initial viral loads (**Figure 2**). In the configuration of initial null viral load (*Q*_*3*_ stock value=0) the value of stocks Q*1*, Q*2A* and Q*2B* were constant and the system behavior was stationary (**Figure 2A**). Assuming a different viral loads (time zero) in the 10-10,000 virions range, we found that there is a threshold in the initial viral load for triggering the progressive reduction of *Q*_*1*_, whose amount could in turn trigger the cell death in different ways. Apoptosis is a cellular process requiring energy, and a deflection in *Q*_*1*,_ as shown in **Figure 2F, Extended Figure 2** and **Supplementary Methods**, suggesting that cell lysis due to energy catastrophe can occur in an infected cell when the baseline viral load is high (range 5,000-10,000 virions). Progressive increase of initial viral load contained in *Q*_*3*_ and *Q*_*4*_ stocks interfers with the processes of the host cell, forcing host-proteins *Q*_*2B*_ to favor the production of virions (progressive increase of the *Q*_*5*_ stock value) through increased *J*_*23*_, *J*_*25*_ and *J*_*27*_ flows (**Figure 2B-F**). Thus, the *Q*_*5*_ dynamic evolution is significantly dependent on the initial configuration (baseline amount of *Q*_*3*_, *Q*_*4*_, *Q*_*2B*_ and cell bioenergetics, represented by *J*_*0*_ and baseline *Q*_*1*_ amount). This could favor virions spread to other cells and changing the system output, with a bulk release of the content of *Q*_*5*_ outside the cell. For a given *Q*_*1*_ basal amount (which represents a specific phase of cell cycle, since *Q*_*1*_ by definition is the amount of ATP-eq allocated for protein synthesis in the cell) and *Q*_*3*_ (equivalent to 10k virions infection), the configuration of the system should be therefore simulated at different time points. The behavior of *Q*_*1*_ following different initial values of *Q*_*3*_ could explain why infections due to hCoVs have a wide incubation time, and in most cases occur asymptomatic, since in half of the possible configurations no relevant changes in the host cell energetics were observed. This phenomenon can have clinical implications, providing a better understanding of the configurations that can favor the action of external driving forces (e.g., drugs).

**Figure 2.**
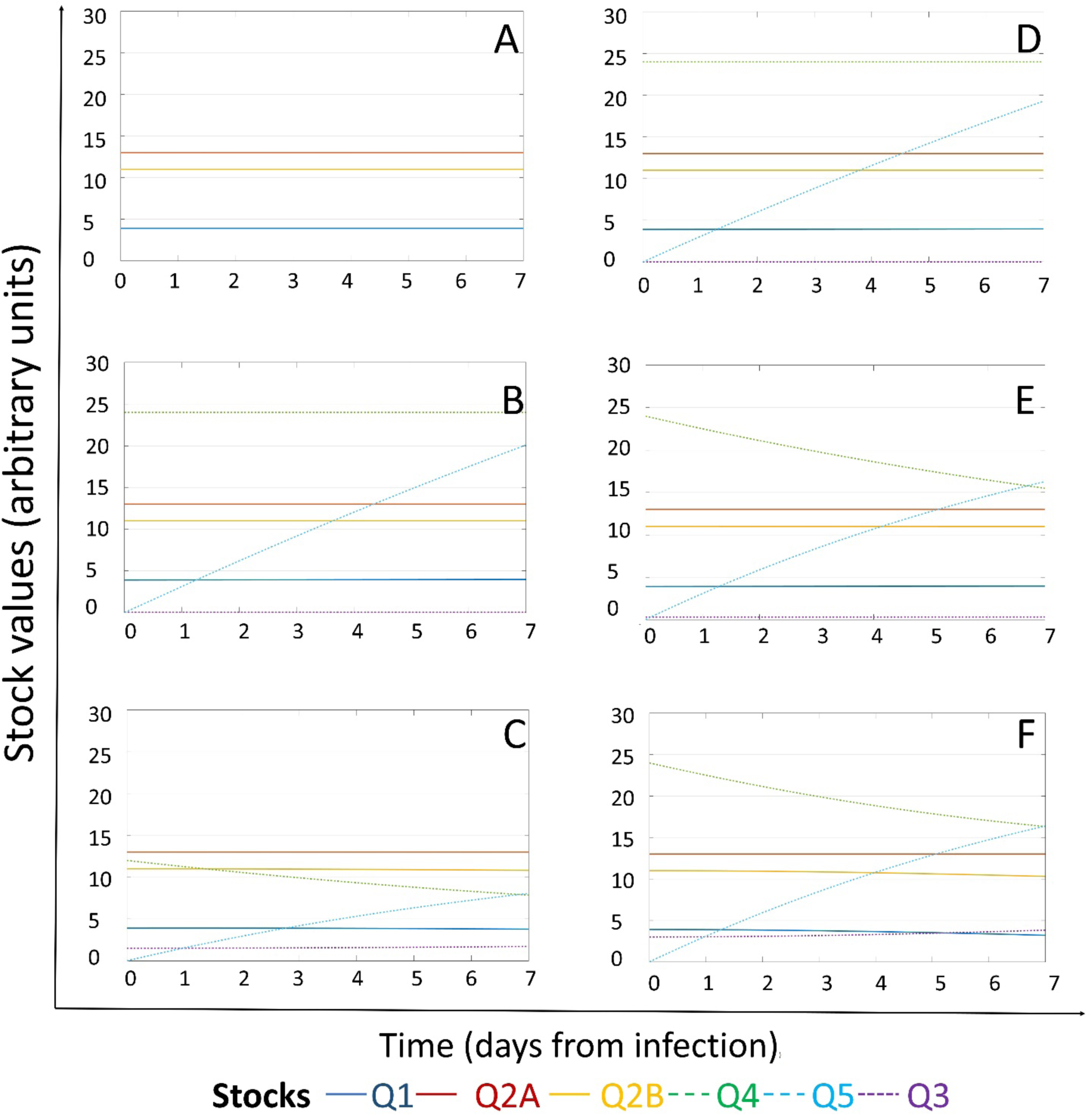
Effects of initial viral load on the energy dynamics of a cell infected by a ss+RNA virus. Stock values, expressed in ATP-eq, are shown from Day 0 Time 0 from infection as a function of different initial viral loads (Initial Q3 values respectively: **A**=0 virions, **B**=10 virions, **C**=100 virions, **D**=1,000 virions, **E**=5,000 virions, **F**=10,000 virions). Color codes for each stock is addressed on the bottom.

### Dynamics of the systems configuration in response to target therapy

When a (+)ssRNA virus enters the single host cell, the *Q*_*3*_ stock is fed, and its proteins can interact with the host proteome to sustain RNA replication. Based on previous works in the field ^59-63^, we identified a time delay of 2 to 6 hours required to record changes in the *Q*_*5*_ stock. Moreover, the value of *Q*_*5*_ varies over the time due to changes occurred at a different timepoints in the stocks *Q*_*2B*_, *Q*_*3*_ and *Q*_*4*_. To be effective, a therapeutic strategy should limit the outflows of virions, *J*_*7*_ and/or *J*_*50*_. However, minimization of *J*_*7*_ seems to be counter-effective in our simulation, due to the increase of *Q*_*5*_ as consequence of the feedback action in the virus replication (VR) process. The minimization of *J*_*50*_ prevents the outflow (viral shedding) from *Q*_*5*_ without stopping its growth. At the same time, this leads to increased resources diverting from *Q*_*1*_ and *Q*_*2B*_ that could promote the cell death, with consequent spread of the virions in the environment.

We applied the search of systemic leverage points by simulations of the diagram dynamics of multiple scenarios. We evaluated several systemic configurations leading to the minimization of the *Q*_*3*_ and/or *Q*_*5*_ stock over time upon the action of a generic external driving force (*D*). We simulated the effects of full (100%) or partial (50%) reduction of either *J*_*3*_ (flow of energy required for RNA replication), or *J*_*4*_ (flow of energy required for viral RNA translation), or *J*_*5*_ (flow of energy required for virions assembly), or *J*_*50*_ (flow of energy required for virions budding), which are typically dependent on intrinsic viral biological properties. The amount of stocks *Q*_*2A*_ and *Q*_*2B*_ remained constant in all simulations, suggesting that the host-cell bioenergetics does not modify protein synthesis, that is used to feed viral protein synthesis as well.

Full (100%) or partial (50%) reduction of *J*_*4*_ and *J*_*50*_ flows at day 0 did not affect the dynamics of the system (**Extended Data Figures 3 A-E, B-F, D-H** and **Figure 3**, respectively), suggesting that these are not good leverage points at this timepoint. On the contrary, external driving forces able to half flows *J*_*3*_ and *J*_*5*_ could induce progressive reduction of *Q*_*3*_, associated to increase in *Q*_*1*_ (**Figure 3**). Full reduction of *J*_*5*_ at Day 0 can recover the systems dynamics in a stationary status (**Extended Data Figure 3 C**), while a partial reduction can only reduce at each timepoint the stock values of *Q*_*4*_ (**Extended Data Figure 3 G**) and *Q*_*5*_ (**Figure 3**). Only reduction in *J*_*5*_ could also reduce *Q*_*5*_ at Day 1, alone or in combination with reduction of *J*_*50*_ and *J*_*21*_. However, the positive effects on *Q*_*1*_, *Q*_*3*_ and *Q*_*5*_ arising from targeting *J*_*5*_ was mitigated by the combination with reduction of *J*_*50*_ or *J*_*21*_ (**Figure 3**). The partial reduction of *J*_*21*_, alone or in combination, did not change significantly the dynamic growth of *Q*_*5*_, induced increase of *Q*_*3*_ at Day 5, though it could prevent a reduction of *Q*_*1*_, to preserve the cell bioenergetics and preventing the cell death by ATP lack.

**Figure 3.**
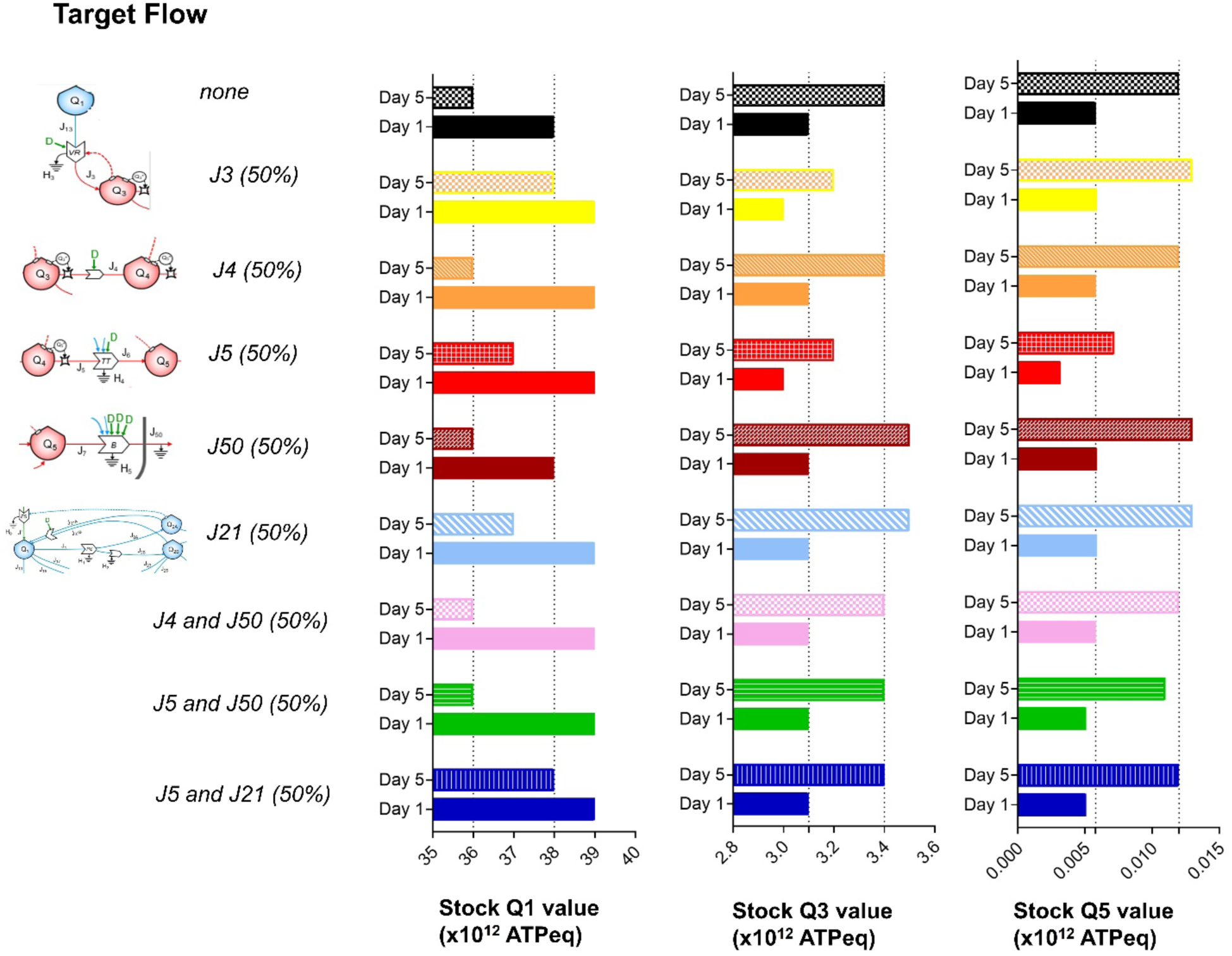
Early and late effects of generic external forcing factors (*F*, e.g.: drugs), at time of infection, on resources available for protein synthesis (Q_1_), viral RNA (Q_3_) and virions (Q_5_) Stocks values *Q*_*1*_, *Q*_*3*_ and *Q*_*5*_, expressed in ATP-eq, were depicted over-time, from infection (Day 0, Time 0) through next 7 days, upon application, at Day 0, Time 0 of generic external forcing factors (*F*) able to reduce the target flows (indicated on the left) of 50%. Values are referred to Day 1 and Day 5 from infection.

### Dynamics of the systems coonfiguration to target therapy given at different times from infection

We also simulated the effect of applying the same external inputs at different times: after 1 (**Extended Data Figure 4**), 3 (**Extended Data Figure 5**) or 5 days (**Extended Data Figure 6**) from the initial infection, alone or in combination (**Figure 4**). Application of external forcing factors at Day 1, Day 3 or Day 5 could affect in a non-linear way the system response, with the largest efficacy targeting *J*_*5*_ (**Figures 4-5**). The combination of external driving forces acting on *J*_*5*_ and *J*_*21*_ did not result to be significantly synergic, while targeting *J*_*50*_ and *J*_*21*_ increased the *Q*_*5*_ stock value instead of the expected reduction. It is worth stressing that this kind of behavior is a typical systemic feature, where an intervention on a specific local process may lead to counterintuitive rearrangements in the system dynamics as a whole. Late suppression of *J*_*5*_ and *J*_*50*_ at Day 5 could reduce only the stock values of *Q*_*4*_. Only early full reduction of *J*_*5*_ (by Day 1) could significantly limit the growth of *Q*_*5*_, while the combination of contemporary suppression of *J*_*5*_ and *J*_*50*_ could not prevent *Q*_*5*_ growth (**Figure 5**). Thus, application of external driving forces at different timepoints is expected to model additional resilience configurations.

**Figure 4.**
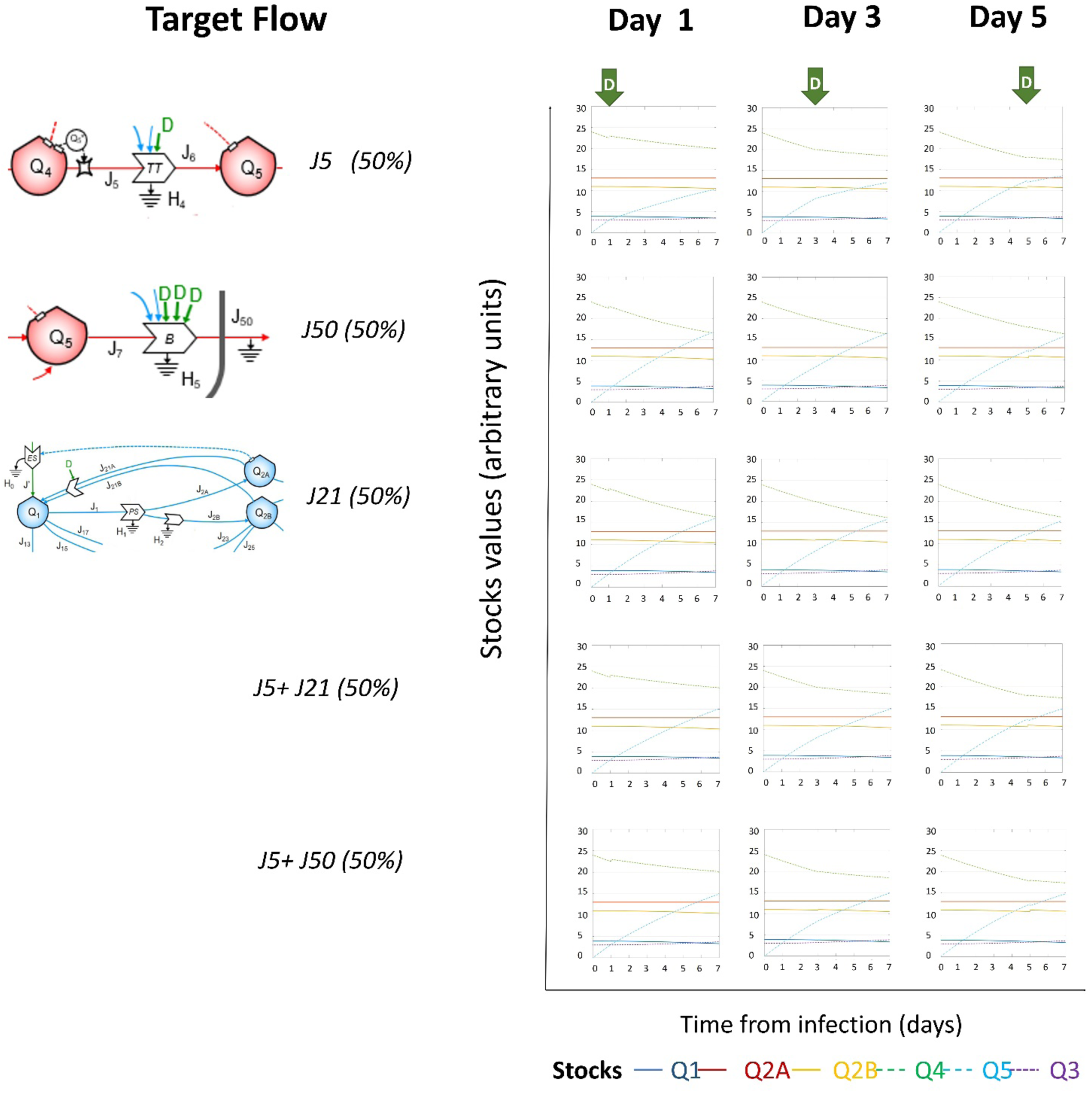
Dynamic changes in system configuration, based on reduction in flows J_5_, J_50_ and J_21_, alone or in combnation, when generic external forcing factors (*F*) are applied at different timepoints (on days +1,+3,+5 from infection) Dynamics of stock values, expressed in ATP-eq, are shown from Day 0 Time 0 from infection as a function of external forcing factors (*F*), able to reduce the target flows (indicated on the left) of 50%, applied at Day 1-3-5. Color codes for each stock is addressed on the bottom.

**Figure 5.**
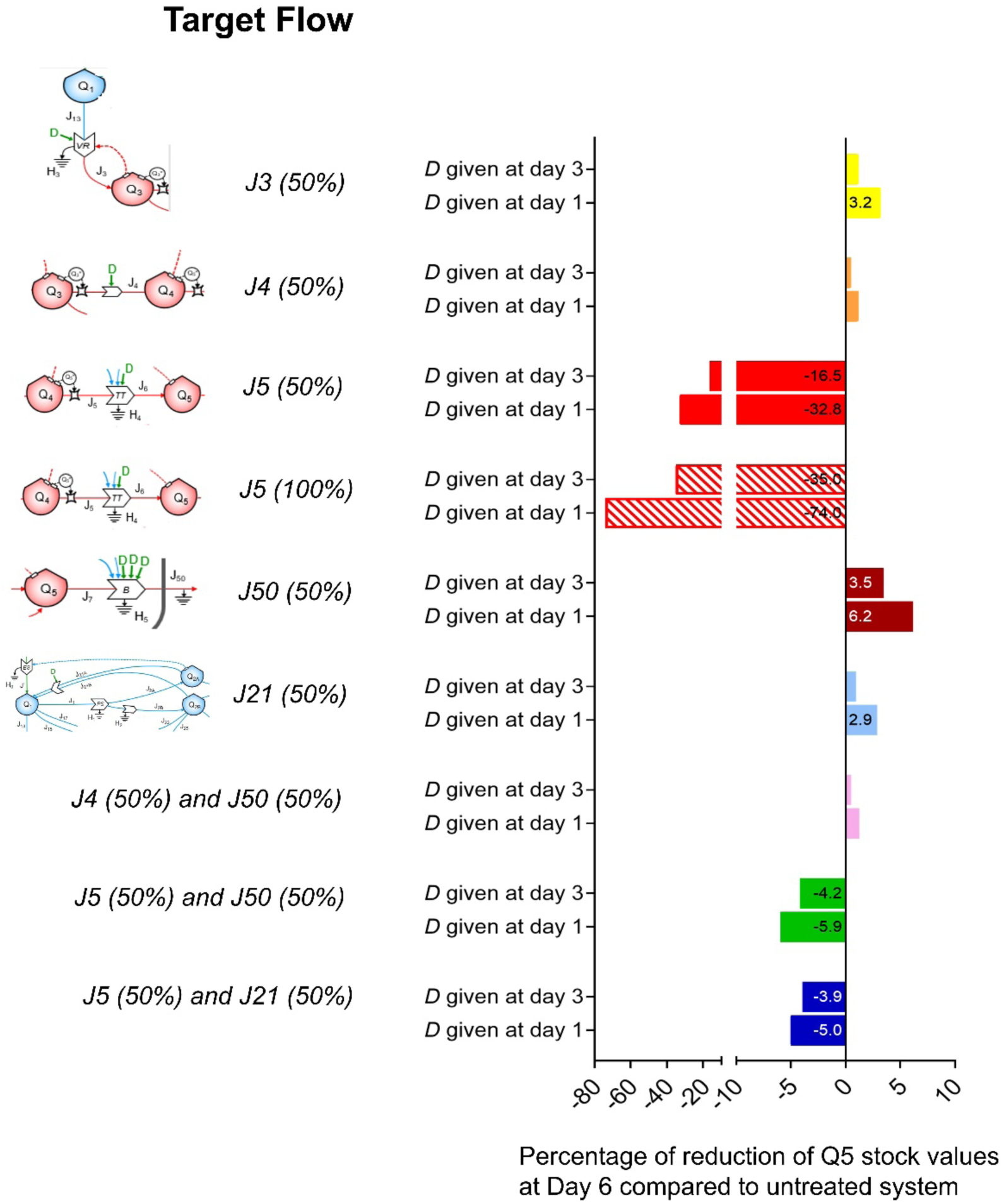
Changes in virion (*Q*_*5*_) values at Day 6, based on external forcing factor (*F*) applied at Day 1 or at Day 3. Percentage of reduction for *Q*_*5*_ stock values at Day 6 compared to Day 0 Time 0 upon application of external forcing factor (*F*) able to reduce the target flows (indicated on the left) of 100-50%, given at Day 1 or at Day 3. Percentage numbers for each simulation are shown in the correspondant bars.

## Discussion

In this work, we approached the host-virus interaction dynamics as a systemic problem and, for the first time in the field, we used combined Systems Thinking tools as a conceptual framework to build up a systemic description of the viral action and host response, critically depending on the existing metabolic environment. The complex dynamic behavior was described in terms of underlying accessible patterns, hierarchical feedback loops, self-organization, and sensitive dependence on external parameters, that were analytically computed by a simulator. The following novelties were addressed: i) the use of energy language as a common quantitative unit for different biological elements in modelling cell-virus dynamics co-evolution; ii) the description of the virus infection dynamics in terms of systemic interaction between virus and host cell stocks, that are included in the same system; iii) the use of a dynamical simulator, that can describe the evolution of cell infection under different conditions; iv) the setting up of a model for different systemic responses to external driving forces (e.g., drugs), that identify different leverage points.

In general, our results address the conceptualization of diseases as complex systems, like in the case of cancer ^64,65^. In fact, the multiple interactions between virus RNA and host cells operate as a dynamic system, capable to overcome the action of external driving forces. This systemic behavior is a major obstacle in the understanding of how developing novel therapies ^66^. Infection works across a range of intertwined spatial and temporal (hours to days) scales ^32^, suggesting that a description able to connect all the levels at which the infection proceeds is needed ^49^.

The presence of a (+)ssRNA-virus stock within the system is one of the possible self-organized patterns (configurations) of a single cell. The virus-host co-evolution ^67^ was represented as an evolving pattern, to which the RNA-virus has given access, using pre-existing stocks, processes and flows of the host cell. Conversely, healthy host-cell dynamics can be represented through a transient stationary state of relevant stocks (i.e., *Q*_*1*_, *Q*_*2A*_ and *Q*_*2B*_). The use of energy as key parameter in modelling the virus-host co-evolution, at the level of the single cell, complies with the medical literature evidence, showing the relevance of energy as a descriptor of biosystems in medicine ^68-71^. Addressing the requirement of a poly-target approach ^72^, our work implemented the description of virus energetics, previously given in terms of predicted energy use in different steps of the viral evolution ^53,73^. Our observations were indirectly confirmed by the data recently published by Gordon et al., who cloned, tagged and expressed 26 of 29 SARS-CoV-2 proteins individually in HEK293T cells and used mass spectrometry to measure protein–protein interactions ^6^. With this experimental approach, they identified 332 interactions between viral and host proteins, and noted 69 existing drugs, known to target host proteins or associated pathways, which interact with SARS-CoV-2, addressing the importance to target the host-virus interaction at the level of RNA translation.

There are known advantages of in silico modelling the action of therapeutic agents on known diseases through agent-based modelling ^74^. However, the literature evidenced some intrinsic limitations on the choice of parameters, like the size of investigated populations ^75^, while major problems are related to model validation ^75,76^, also requiring to supplement the models with adequate formal ontologies ^77^. Thanks to its abstract nature, stock-flow description can be used in a wide range of different fields, realizing the conceptual bridge that connects the language of biological systems to that of ecology.

Our approach unveils the potential of systems thinking for the study of other diseases or classes of disease, since it appears more and more clear how some incurable pathologies can be described only by adopting a more comprehensive systemic approach, in which the network of relationships between biological elements are treated in quantitative way similar to that applied in this paper.

## Methods

The method used to study the dynamics of the virus-host interaction system is structured in 3 basic steps, namely, the development of a flow-stock diagram, that describes the virus-host interactions, the development of a virus-host systemic simulator, and, finally, its calibration and validation (**Extended Data, Figure 1**).

### Development of the stock-flow diagram

A typical Systems Thinking diagram is formed of stocks, flows and processes. Stocks are *countable extensive variables Q*_*i*_, *i*=1,2,…,*n*, relevant to the study at issue, that constitute an n-ple of numbers that at any time represents *a state of the system*. A stock may change its value only upon its inflows and/or its outflows, represented by arrows entering or exiting the stock. Processes are any occurrence capable to alter – either quantitatively or qualitatively – a flow, by the action of one or more of the system elements. In a stationary state of the system, stocks values are either constant or regularly oscillating. In the dynamics of a system, stocks act as shock absorbers, buffers, and time-delayers. Processes are all what happens inside a system that allows the stationarity of its state, or that may perturb the state itself. To occur, a process must be activated by another driver, acting on the flow where the process is located. These interaction flows may be regarded as flows of information, that control the occurring processes and so the value and nature of the matter flows. The pattern of the feedbacks acting in the system configurations is the feature that utlimately defines the systems dynamics.

### Development of the systemic simulator

We adopted an energyy approach, where stocks, flows and processes are expressed in terms of the energy embedded, transmitted and used, respectively, during the system operation. The equations, that characterize the flows relevant to the diagram, are typical of dynamic ST analysis ^51,78^, and their setting up is linked in many respects to the energy network language ^79^. In this approach, each flow depends on the state variables *Q*_*i*_ by relationships of the kind *dQ*_*i*_*/dt=kf(Q*_*j*_*), i,j*=1,…,*n*, where *n* is the number of stocks in the system. Given a set of proper initial conditions for the stocks (i.e., the initial system state) and a properly chosen set of phenomenological coefficients *k*, the set of interconnected equations will be treated by standard finite-different method, taking care of choosing a time-step short enough to evidence the possible dynamics of any of the studied processes. The coefficients *k*_*i*_ are calculated using data on the dynamics of any single stock, in particular, by estimating the flows and the stocks during the time interval set for the simulation steps (as described in **Supplementary Methods**). When different flows co-participate in a process, a single coefficient will gather all the actions that concur to the intensity of the outcoming flow(s). These phenomenological coefficients are not, therefore, related to specific biophysical phenomena or processes, but are set to describe how and how fast any part of the system react to a change in any of the other ones. In general, our model may be then regarded as based on a population-level model (PLM) as defined by ^80^, which can be run at different scales from organism to sub-cellular ones (see for example ^58,81,82^), that already increased the existing knowledge on the life cycle of different infections, like for example the ones generated by HIV and Hepatitis C virus ^83^. To obtain the simulations, we used the open-source computation software SCILAB (https://www.scilab.org).

### Calibration and validation of the simulator

The model simulations are developed based on the choice of the initial stock conditions, as well as of the parameters *k*_*i*_ (see **Supplementary Information** for further details). A model is valid as far as its output reproduces the reality, and systems modelling must be tested under two aspects, concerned with the capability to correctly describe the system and its dynamics, respectively ^84^. The former aspect has to do with the completeness of the choice of stocks and of inflows-outflows structure. The latter is related to the completeness and the correctness of the relationships between the stocks, i.e., the setting up of the equations that represent the matter flows as well as the control actions that the stocks exert on processes. A single comprehensive procedure for the of validation of all dynamic models however does not exist ^85^, since the validity of a well-made model is anyway dependent also on its usefulness, referred to the very purpose of the model itself. Knowledge in the field must guide the setting up of the model, in order to legitimate the choice of the elements that form the systemic simulator (see **Supplementary Materials** for further details). This was done using both information from existing knowledge on the biological mechanisms at issue, and experiment-based calculations, that allow to the identification of the variables and parameters necessary to set up the equations describing the system dynamics. This evaluation procedure encompasses both the static features and the dynamics of the system. In our case, the first fundamental test was the verification of the prediction of a stationary state for the values of the stocks of the system in the absence of contagion, i.e., when no competition is occurring between virus and the host. The model itself was set up by an iterative process, in which the scientific medical knowledge about the virus was used to draw a first model, then converted to a simulator, refined and integrated by testing its ability to simulate real data and the actual evolution of the disease under different conditions. The reliability of both available data and modelling were tested by evaluating the variation of the model output upon “artificial” changes of each of the most relevant input data. Some of these numbers are affected by relatively high uncertainties. Thus, a sensitivity analysis was performed on the outcome of the simulations, testing to what extent a possible variation on these values lead to incorrect evaluation of the system dynamics. For this reason, we applied a 50% variation (either positive or negative), in different combinations, to the parameters on which the results are more sensitive. As expected, the corresponding simulations vary as well, but the general dynamics of any of them resulted unchanged, especially concerning the overall trends shown by comparing the groups of simulation, as in the figures presented in this work.

## Data availability

The inventory of initial values of stocks and phenomenological coefficients K are available in **Supplementary Methods Tables 1** and **2**. The code is available upon request and described in the **Supplementary Methods**.

## Acknowledgements

This work was supported by A.I.L. (Associazione Italiana contro le Leucemie) sezione di Catania and FON.CA.NE.SA. (Fondazione Catanese per lo Studio delle Malattie Neoplastiche del Sangue).

## Author contributions

A.R. designed the study and collected medical knowledge, F.G. built up the diagram and M.C. built up the simulator. All authors performed simulation, analysed data, prepared the figures and wrote the paper.

## Extended data

**Extended Data Figure 1.**
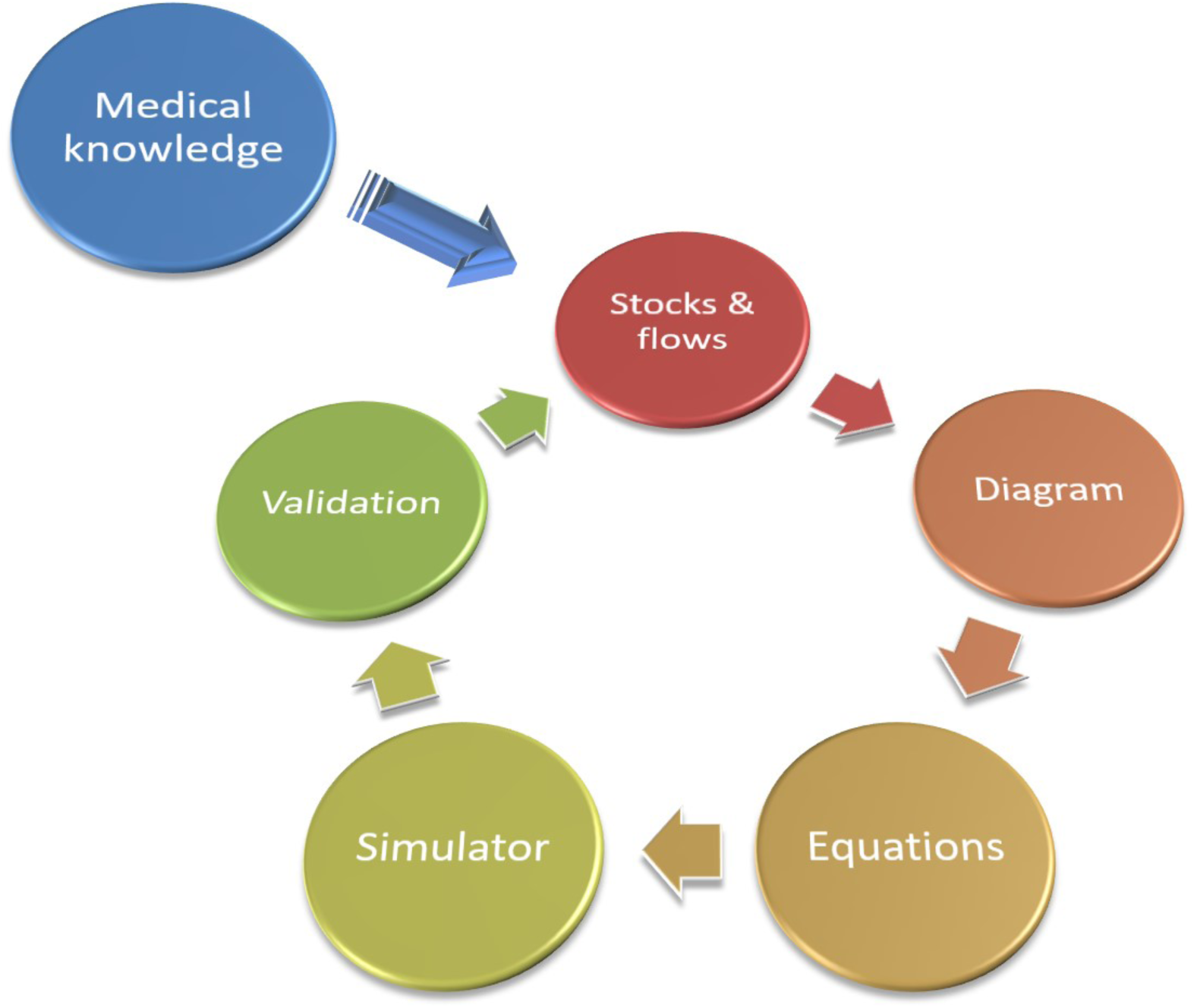
Workflow of study design. In order to set up suitable diagram simulators and so to operate quantitative analyses, a proper knowledge of the biology of the system is necessary, along with reliable and comprehensive data sets (medical knowledge). Once defined the boundary conditions, stocks, flows and processes are included in the ST diagram. An inventory including the initial values, expressed in the same unit of measure, and the phenomenological coefficients required for the differential equations that make the simulator runs. Resulting simulations are compared with the experimental observations and validated by knowledge in the field.

**Extended Data Figure 2.**
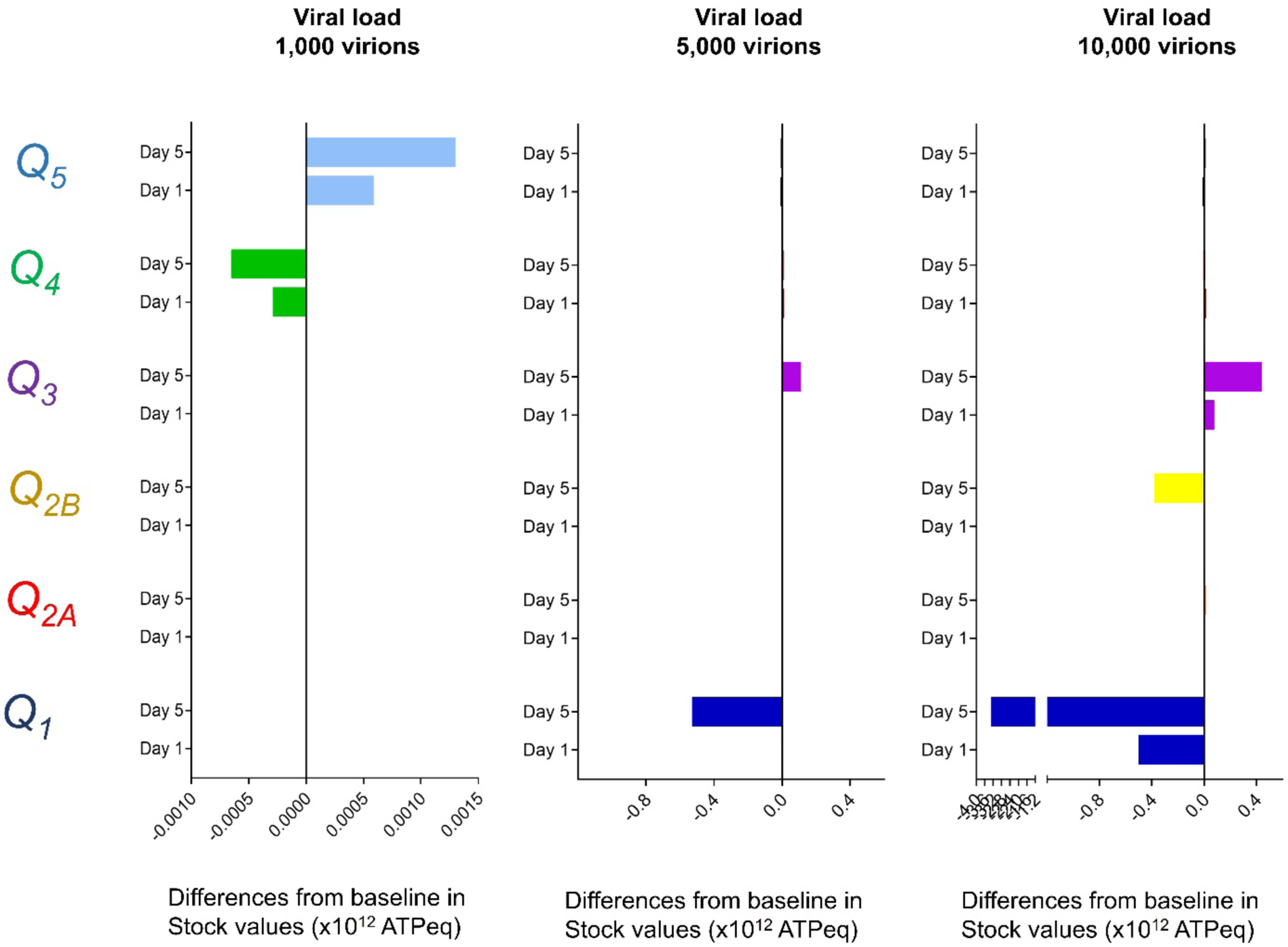
Early and late changes in stocks values based on viral load at Day 0. Time change of the stocks values, expressed in ATP-eq, from infection (Day 0, Time 0) through the next 7 days, using different initial values in *Q*_*3*_ (amount of 1,000-5,000-10,000 virions, expressed in ATP-eq). The difference between the value at Day 0, Time 0 and that after 48 hours (at the end of Day 1) or 144 hours (at the end of Day 5) is plotted for each stock.

**Extended Data Figure 3.**
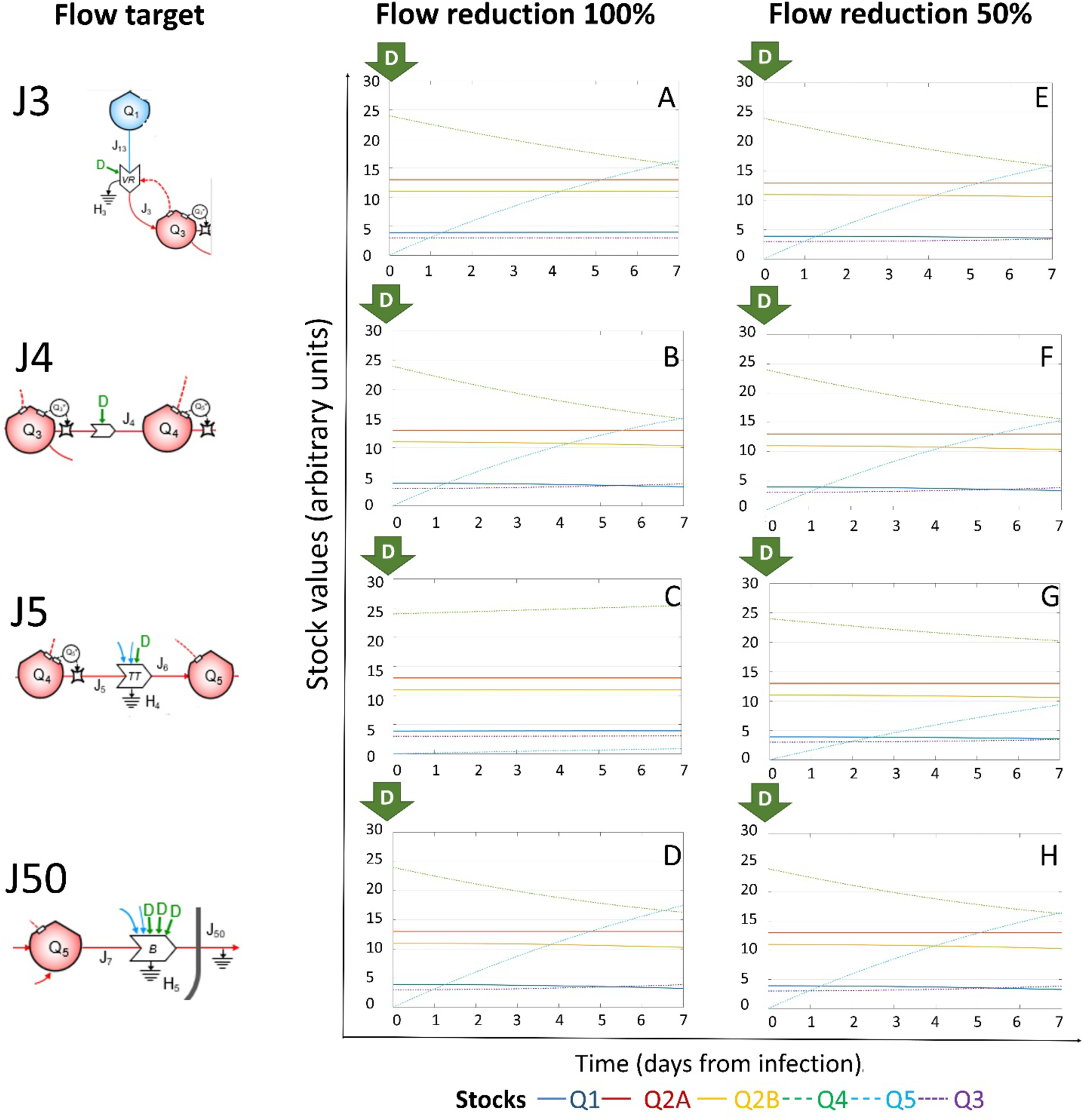
Effects of targeting leverage points applying generic external driving force (*D*) at Day 0. Time change of the stocks values (for the color code see at the bottom), expressed in ATP-eq, from infection (Day 0, Time 0) through the next 7 days, upon application, at Day 0, Time 0 of generic external driving forces (*D*) able to reduce the target flows (schemed on the left) of 100% or 50%, respectively.

**Extended Data Figure 4.**
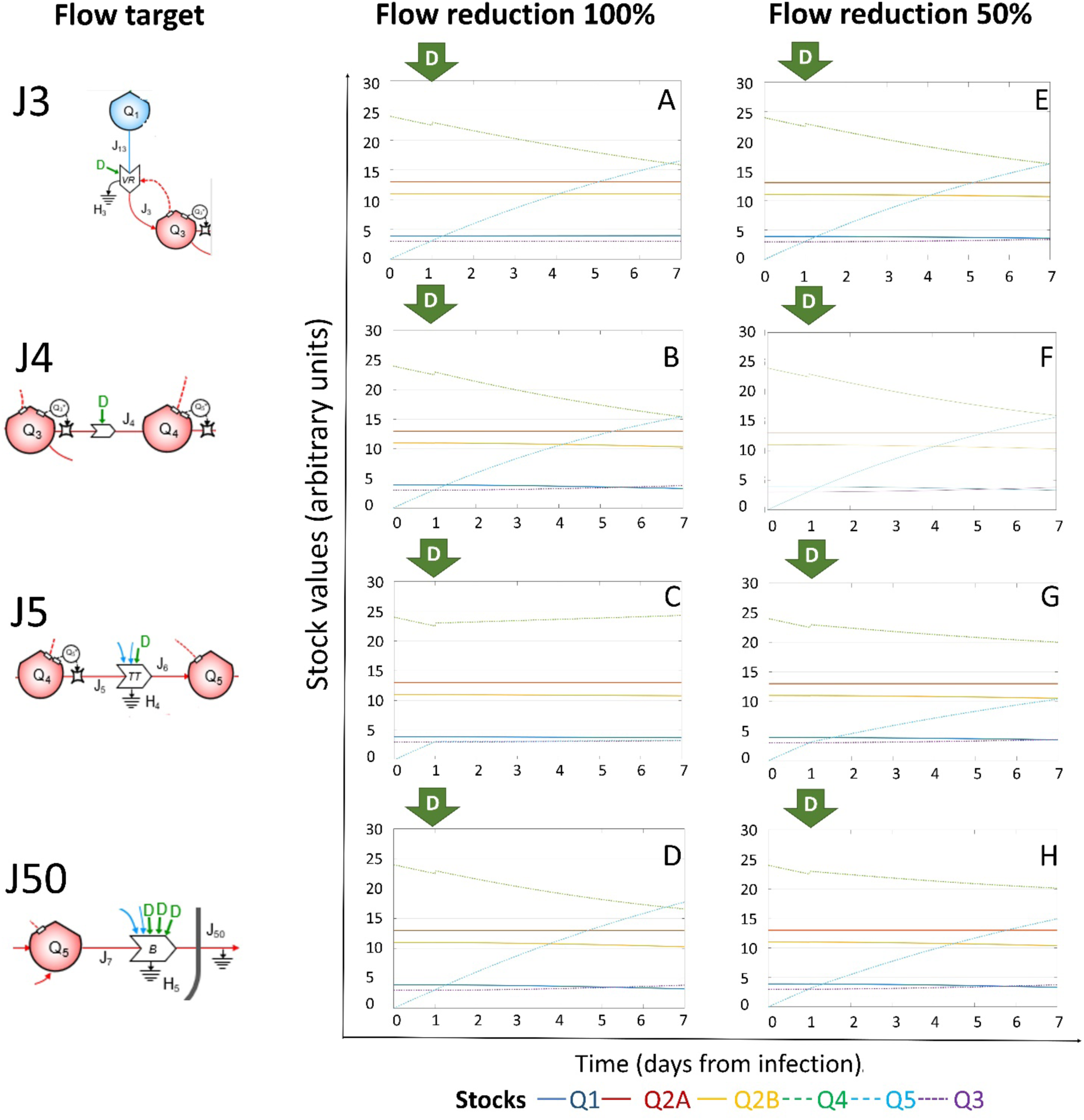
Effects of targeting leverage points applying generic external driving force (*D*) at Day 1. Time change of the stocks values (for the color code see at the bottom), expressed in ATP-eq, from infection (Day 0, Time 0) through the next 7 days, upon application, at Day 1, of generic external driving forces (*D*) able to reduce the target flows (schemed on the left) of 100% or 50%, respectively.

**Extended Data Figure 5.**
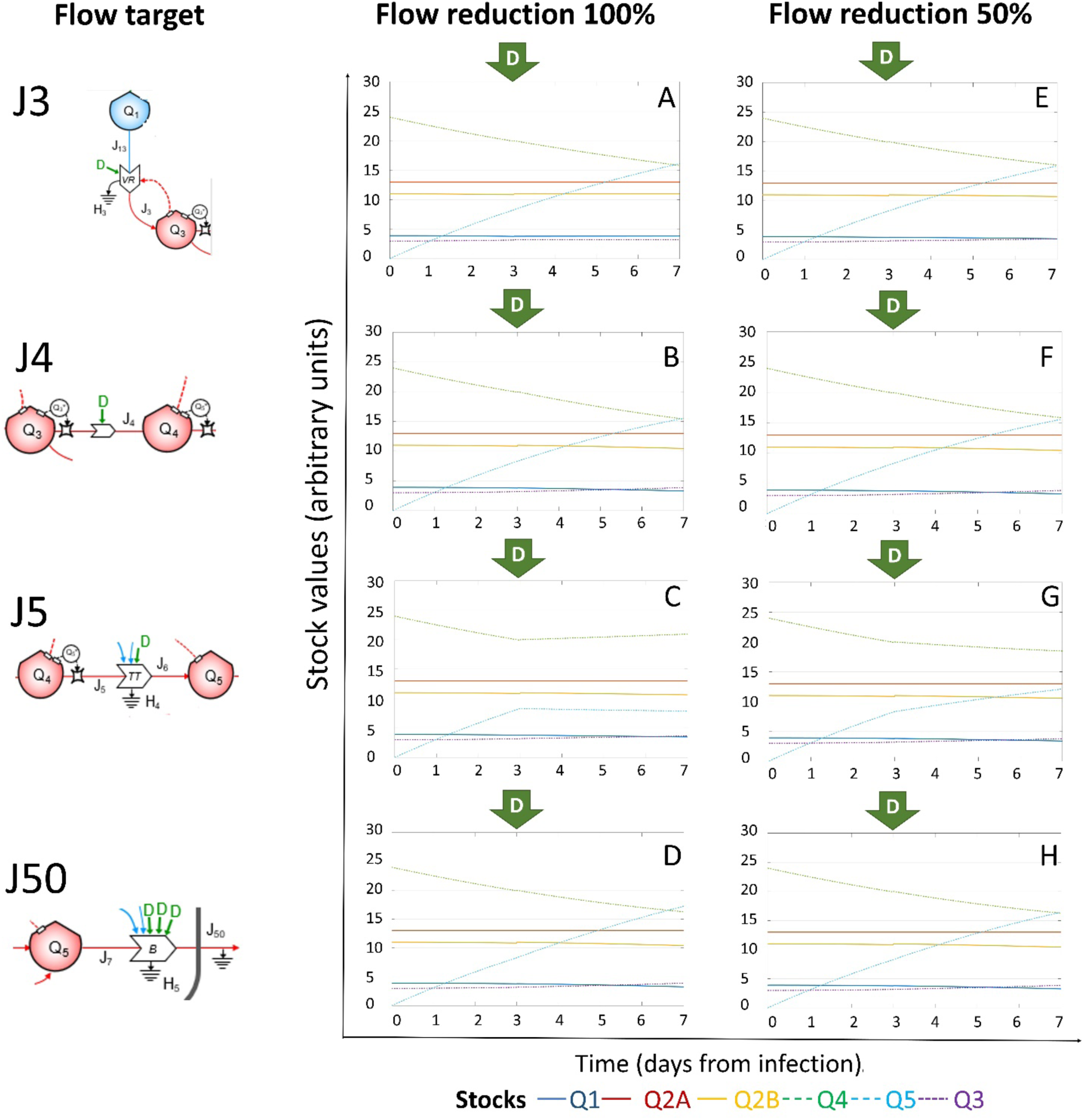
Effects of targeting leverage point applying external driving force (*D*) at Day 3. Time change of the stocks values (for the color code see at the bottom), expressed in ATP-eq, were depicted over-time, from infection (Day 0, Time 0) through the next 7 days, upon application, at Day 3, of generic external driving forces (*D*) able to reduce the target flows (schemed on the left) of 100% or 50%, respectively.

**Extended Data Figure 6.**
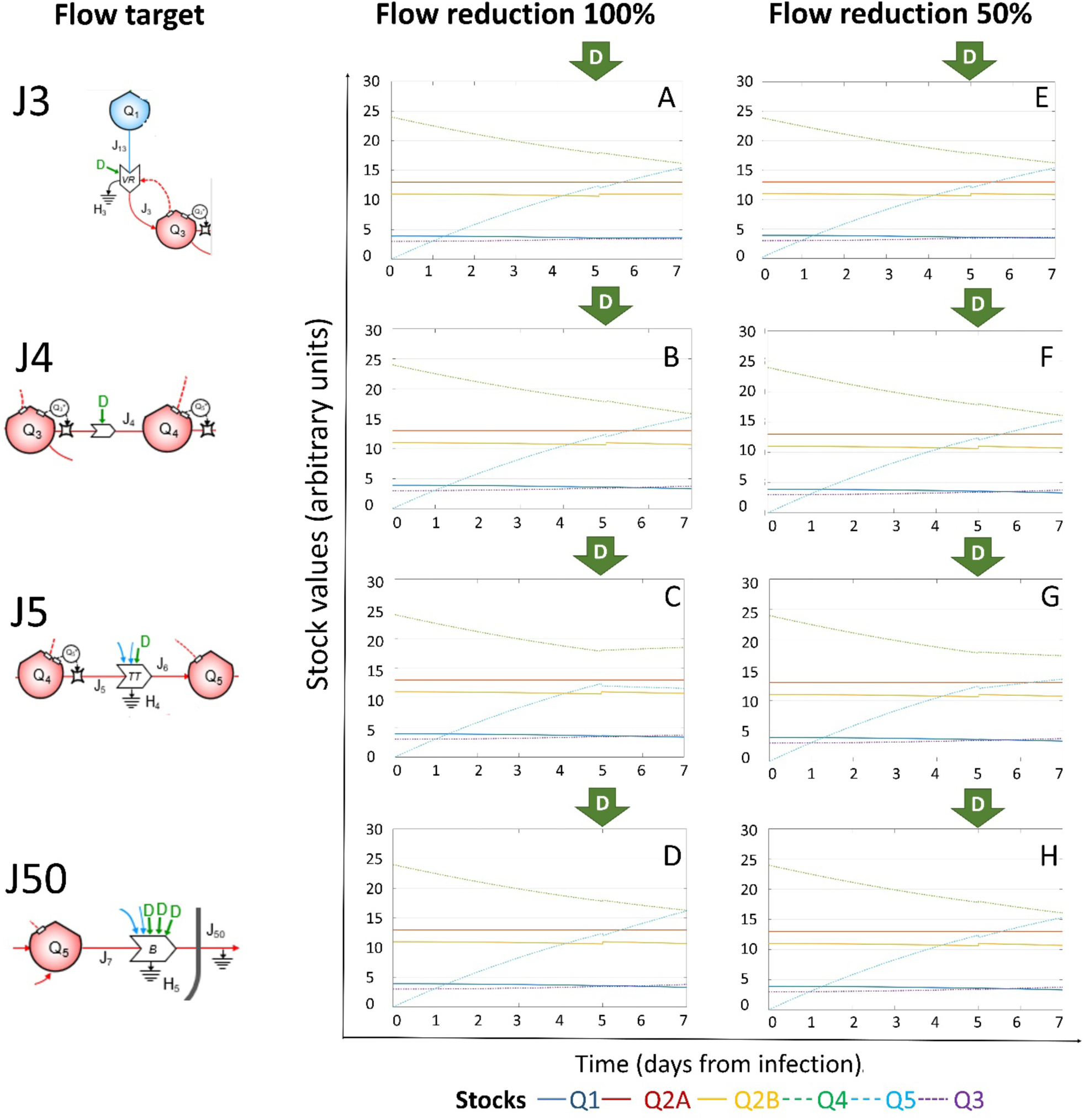
Effects of targeting leverage point applying external driving force (*D*) at Day 5. Time change of the stocks values (for the color code see at the bottom), expressed in ATP-eq, were depicted over-time, from infection (Day 0, Time 0) through the next 7 days, upon application, at Day 5, of generic external driving forces (*D*) able to reduce the target flows (schemed on the left) of 100% or 50%, respectively.

