## Supplementary information for "Energy dynamics for systemic configurations of virus-host co-evolution"

**Supplementary Discussion**

In this study, we captured the dynamic co-evolution of host-cell bioenergetics in presence of a ss+RNA virus, using a Systems Thinking (ST) approach.

_Advantages and limitations of ST approach_

In systems thinking, the behavior of a dynamic system is not described from a reductionist point of view, by specifying the operation of any of its elements, nor by statistical processing of big data-based networks. Instead, the study is focused in the underlying systemic patterns, hierarchical feedback loops and self-organization, and can be analytically computed by proper simulators [1-4], that may therefore address suitable leverage points for intervening [5]. The basic ST idea is to integrate the traditional bottom-up approach – that describes "*local*" behaviors through cause-effect chains and functional units – with a *top-down* approach, that points out at the global behavior of the system in terms of its operational configurations, emerging from the feedback-driven response to different driving forces, like those represented for example by the chemistry of new drugs [6]. Systems Biology already recognized the relevance of complexity in the study of microbiological systems [7,8], but though successfully applied in several fields ranging from hard sciences to ecology and economics [9,10], the potential of ST in the study of biological systems is still underexploited. Systemic approaches have started catching on in biology and medicine in the last decades mostly as computational-based tools, owing to the development of network analysis (NA) procedures and the concurrent availability of large set of data. Concepts like system biology [7,8] [4,11], bioinformatics [12], network medicine [13,14], network pharmacology [15,16], machine and deep learning [17,18] are now used in many fields of cancer research, often regarding the disease as some kind of network perturbation [19] to be investigated by computational means [3,20]. Contrary to expectations, and despite the efforts devoted to the NA of genes, proteins and metabolites, network approaches have not yet delivered the expected revolution for instance in cancer comprehension [21] due the enormous non-genetic plasticity of tumor cells [22], the ambiguity in determining whether the specific biological effects depend on individual or simultaneous modulations of established targets [23] or information loss depending on network structure initial configuration [16].

In this framework, the search for a therapeutic strategy is usually based on target-related parameters. In our case, we have shifted the attention from target-related to systemic parameters, as suggested also by recent literature [24]. ST approach starts from the identification of a limited number of state variables, those necessary to model the resource flows and the feedback networks that describe the configuration patterns of the system dynamics. Indeed, offering a novel perspective [25-27], the complementarity between ST and NA presents itself an enormous potential, to accelerate disease evolution comprehension [28]. Though sharing a general systemic perspective, a fundamental difference however stands between ST and NA in terms of the system elements. In NA, the elements are vertices representing the entities whose network is called to describe the system at issue. They are connected each other by edges, that may take the form of exchanges of something, physical connections (like the roads in a transport network), neuron synapses, friendship relationships, paper citations, predation of species, and so on. They are required only to represent an existing interaction between the vertices, to form a network for which big data are or may be available. In the ST description, stocks and flows refer to measurable, physically identifiable resources (matter, energy, information) and they are not a list of systems elements, like nodes and edges used in the network analysis. In a sense, these latter consider a system as the collection of many elementary elements, whose dynamics is investigated thanks to the availability of big data and application of mathematical and statistical tools [13,29-32].

Systems Thinking is, therefore, both a conceptual framework and an analytical tool for the study of complex systems. ST approach starts from the identification of a limited number of state variables, the stocks, those necessary to model the resource flows and the feedback networks that describe the configuration patterns of the system dynamics. The conceptual relevance and the potential of Systems Thinking approaches – in particular, the use of stock-flow representations – lie in the holistic top-down description of systems, whose complexity and dynamics can be hardly represented by reductionist, bottom up approaches. This is due to the sensitivity of the overall systemic behavior to the configuration feedbacks network, connecting the elements of the system. Depending on the hierarchical level chosen for the study (in our case, the cell level), a different description of the system will be given in terms of stocks, flows and processes. Consequently, in order to set up suitable diagram simulators and so to operate quantitative analyses, a proper knowledge of the biology of the system is necessary, along with reliable and comprehensive data sets. They are necessary to define the boundary conditions, the initial values and the phenomenological coefficients entering the differential equations that make the simulator run. When the ST analysis is downscaled to the microbiology domain, big data-based systemic approaches become anyway relevant. We think that the future medical research will find it the convergence of ST-based and big-data based network analysis methods an important room for a methodological revolution, that will reconcile in a synergic way the two mindsets.

*Energetics of virus-host co-evolution*

The virus-host co-evolution is based on a common pool of resources, whose dynamics became the first target of our simulations. The availability of recent literature studies on virus energetics at single-cell level [33] constituted the basis for the comparability of variables used to quantify the system dynamics. The search for detailed molecular mechanisms, by which specific viruses target their replication factors and their RNAs to specific membranes or intracellular sites, to assemble replication complexes or factories, still remain unaddressed for several ss+RNA viruses [34]. Similarly, it should be investigated how different viruses coordinate the various and often complex membrane rearrangements associated with their replication processes [35]. Moreover, the assemblage mechanism of RNA replication complexes on distinct secretory, endosomal, or organellar membranes, and the implications of such choices for replicative efficiency, virus-host interactions, and pathology for +strand RNA viruses remains as well poorly understood.

*Study limitation and future perspectives*

The model proposed in this work was developed at single-cell scale. However, in order to define an overall therapeutic approach, a multi-scale approach would be also desirable. In particular, depending on the availability of appropriate data, a future model could focus on different scales, with a more detailed description of some components at sub-cellular level, that were grouped (e.g., short and long half-life proteins, lipids and vesicles trafficking) in the present study. On the other hand, the interaction between different cell populations in the host should be also developed, to represent the interaction between healthy and infected cells, and the contribution of immune system [36-38] or the repertoire of receptors on the surface of the host cell [35,39,40] to surveil and limit the size of $Q_3$ stock at single cells level. Other natural system constraints could be included, like some physiological parameters (e.g. temperature, metabolic rate), whose impact on the human body energy dynamics is already understood. The use of a multi-scale hierarchical perspective is already in principle possible, as discussed in previous works adopting the same sort of system representation [41,42].
Thus, instead of targeting a single biochemical process, the proposed simulations allowed to apply a poly-target logic, in order to identify the most vulnerable processes of viral growth inside a cell from an energy perspective. In particular, results allowed us to move from a complementary perspective with respect to the search of pharmacological molecules, searching for pharmacological poly-targets, that were identified accordingly. It is worth underlining that, by the proposed simulator, a wide range of computational combinations of the different driving forces may be investigated. If these driving forces represent for instance the administration of drugs, simulations can be performed by varying their relative strength, the therapeutic target, the order and the time of administration, and the combination of different ones at the same time. Furthermore, in the overall study of a disease behavior, the plasticity of the approach and of the relative simulator can become crucial when a driving force is the result of another hierarchical level of the same system. This is the case for example of the amount of virus genome mutation, possibly occurring outside the cell but within the super-system that supports the cell functioning. In this case, the accumulation of virus mutations affects the value of internal stocks, e.g., $Q_3$, and thus of the related flows. In this work, we have performed preliminary simulations for a wide range of parameter combinations, and -based on the results- we have presented some of the most representative ones in terms of the method efficacy.

*Future clinical implications*

The advantage of using an ST-based approach is reflected in the possibility of extracting also systemic dynamic features, that would be otherwise counterintuitive. While a traditional single target approach would address strategies targeting the viral RNA ($Q_3$) or the replication process ($J_3$), our results suggest that the virus growth is more vulnerable, if the virion growth before expulsion (process TT, involving flow $J_5$) is targeted. For example, from the start of the COVID-19 outbreak, medical practitioners have followed China's guidelines set up [43] in January and treated hospitalized patients with α-interferon combined with the repurposed drug Kaletra [44-46], an approved cocktail of the HIV protease inhibitors ritonavir and lopinavir, while several trials are ongoing to test the safety and efficacy of hydroxychloroquine alone or in combination with macrolides [47-54] or antiviral Remdesivir, given the synergism in vitro [55]. Waiting for the results of ongoing randomized, controlled clinical trials, Remdesivir showed limited clinical efficacy [56]. Other trials are ongoing to test the optimal treatment and contain the outbreak.

There is an emerging need of tools that could early identify those compounds, not primarily designed for their antiviral action, identifiable by *in-silico* approaches [34], that alone or in combination can provide clinical efficacy [34,57-60]. However, pressing questions remain about how to accelerate randomized clinical trials and avoid unnecessary duplication of efforts [61].

Looking at the pharmacodynamics of compounds currently under investigation [61], we could re-classify them, based on their systemic mechanisms of action (**Supplementary Discussion Table 1**). Their effects could be potentially simulated to establish the single-cell effect, the best time and/or schedule of administration, as shot-cut of in-vitro studies, with a detail level established on the basis of the purpose of the study design. For example, in our study, we described the effects of both chloroquine and hydroxychloroquine, that can interfere with host-cell proteostasis ($J_{21}$ inhibition) and virion assembly ($J_5$ inhibition). Similarly, we studied the impact of antiviral agents such as lopinavir, that mainly interfere with RNA synthesis ($J_3$ inhibition) and consequently on virion assembly.

### Supplementary Discussion Table 1

### Examples of drugs that could act as external driving forces on identified systemic flow targets

| Flow target(s) | Compound | Mechanism of action | References |
|---|---|---|---|
| $J_0$, $J_3$, $J_4$, $J_{21A}$, $J_{21B}$ | FK506 (tacrolimus) | FKBP15 inhibitor<br>ER protein quality control regulators<br>Bioenergetics regulators<br>mRNA translation inhibitor | 34,62 |
| $J_0$, $J_4$, $J_{21A}$, $J_{21B}$ | Rapamycin | | 34,63,64 |
| $J_1$,$J_3$, $J_{21B}$ | Valproic acid | HDAC2 inhibitor | 34,65-67 |
| $J_1$, $J_4$ | SAHA | pan HDAC inhibitors | 68 |
| $J_1$, $J_{35}$, $J_{2B}$ | Selinexor | mRNA nuclear export complex inhibitor | 34,69-73 |
| $J_4$, $J_{2A}$, $J_{2B}$ | Dabrafenib | Kinase inhibitor, protein synthesis inhibitor | 34,74,75 |
| $J_0$, $J_{13}$ | Metformin | Inhibitor of respiratory electron transport, glycolysis regulation | 34,76 |
| $J_3$, $J_{13}$ | Camostat<br>Nafamostat | TMPRSS inhibitors proteolytic cleavage of viral spike protein priming to the receptor ACE2 present in human cell | 34,40 |
| $J_4$, $J_{2A}$, $J_{2B}$ | Ponatinib | Kinase inhibitor, protein synthesis inhibitor | 34,74 |
| $J_3$ | Ribavirin | Nucleoside inhibitor (mutagenic ribonucleoside) | 34,77,78 |
| $J_{13}$ | Chloramphenicol, Tigecycline, Linezolid | Antibiotics, able to inhibit mitochondrial ribosomes | 34,79 |
| $J_{21A}$, $J_{21B}$ | Chloroquine | SIGMAR1/SIGMAR2 inhibitor<br>Autophagy inhibitor | 34,80-84 |
| $J_3$, $J_5$, $J_{21A}$, $J_{21B}$ | Hydrossychloroquine | Autophagy inhibitor, antiviral effect | 34,51,52,80 |
| $J_1$, $J_4$, $J_5$ | Zotatifin, ternatin 4, tomvosertib | mRNA translation inhibitors | 34 |
| $J_1$, $J_{2B}$, $J_{23}$ | Silmitasertib or TMCB | Casein kinase II inhibitors, protein synthesis inhibitor | 34,85,86 |
| $J_3$ | Remdesivir | Nucleoside analog, interfers with RNA-dependent RNA polymerase | 87 |
| $J_{13}$ | Umifenovir | Inhibitor of the fusion between the viral envelope (surrounding the viral capsid) and the cell membrane of the target cell | 88,89 |
| $J_{13}$ | Lisinopril, losartan | ACE inhibitors, prevent the fusion between the viral envelope and the cell membrane of the target cell | 35 |
| $J_3$ | Favipiravir | Selective inhibitor of RNA-dependent RNA polymerase | 90,91 |
| $J_{2A}$, $J_{2B}$, $J_{3*}$, $J_4$, $J_{7*}$, $J_{21}$ | Macrolide antibiotics<br>Azythromicin*,<br>Clarithromycin | Inhibition of ribosomal translation<br>Autophagy inhibition | 92-95 |

### Supplementary Methods

A typical Systems Thinking diagram is formed of stocks, flows and processes. Processes are any occurrence capable to alter – either quantitatively or qualitatively – a flow, by the action of one or more of the system elements. Stocks are countable extensive variables $Q_i$, $i$=1,2,...,n, relevant to the study at issue, that constitute an n-ple of numbers that at any time represents a state of the system. The choice of the set of variables depends on the hierarchical level of the desired description as well as on the overall purpose of the study. The choice of the stocks is crucial for an effective systemic description, at the same time addressing the disruptive power and the plasticity of the approach. Stocks must be chosen respecting some requirements, namely: i) the number of the stocks must be the minimum necessary to describe the state of the system for the prescribed purposes; ii) it must be possible to describe any relevant process occurring in the system in terms of stocks interactions; iii) any system change (either detectable from the external or not) must correspond to a change in the n-ple of state variables; iv) all the stocks should have – in principle – measurables values.

It must be stressed that the flows do not represent some kind of generic interaction (edge), or some logical nexus, like in network analysis, but rather physical flows able to alter the value of a stock. If this change in the stock alters as well the value of the flow which alters the stock, and so on in a cause-effect loop, we say we are in presence of a feedback. This may be direct or indirect, the latter when the reciprocal change of flow and stock values is due to a path that includes other stocks. The pattern of feedbacks is the feature that defines the systems dynamics. The need for a correct representation of all the feedbacks relevant for the system dynamics description will determine also the best choice for the set of stocks.

System operation was modelled through a set of the differential equations, describing the rate of change of the single stocks, and used to set-up a computational simulator. Then, a diagram displaying the system structure in terms of system observables (stocks), flows and processes has been set up.

The complexity of the system prevents an easy straightforward attribution of the respective role of these three variables in the stock supply, because of the network of feedbacks linking all the stocks that ultimately define the dynamics of the system.

Key processes and turnover times relevant for the RNA-virus-host interactions [29,96-127] were derived from the available literature (as summarized in **Supplementary Methods Table 1**).

**Supplementary Methods Table 1**

Inventory of stocks depicted in the diagram of Figure 1.

| Stock | Biological role | Dynamic equation | Initial value | Reference |
|---|---|---|---|---|
| $Q_1$ | Resources available for protein synthesis | $dQ_1/dt = J_0 + J_{21A} + J_{21B} - J_1 - J_{13} - J_{15} - J_{17}$ | 3.9 | 1,6,128 |
| $Q_{2A}$ | Short-half-life protein | $dQ_{2A}/dt = J_{2A} - J_{21A} - J_{20A}$ | 13 | 129,130,131 |
| $Q_{2B}$ | Long-half-life proteins | $dQ_{2B}/dt = J_{2B} - J_{21B} - J_{23} - J_{25} - J_{27} - J_{20B}$ | 13 | 129,130,131 |
| $Q_3$ | Viral ss+RNA | $dQ_3/dt = J_3 - J_4$ | 3 | 132 |
| $Q_4$ | Viral proteins | $dQ_4/dt = J_4 - J_5$ | 0.024 | 133 |
| $Q_5$ | Virions | $dQ_5/dt = J_6 - J_7$ | 0 | 33 |

| Energy flow and Mathematical expression | Involved biological process | Phenomenological coefficient (k) | Reference |
|---|---|---|---|
| $J_0 = k_0 \times R \times (1+Q_{2A})$ | Enter of resources allocated for protein synthesis | 3.9E–06 | 6 |
| $J_1 = k_1 \times Q_1$ | Host-cell RNA transcription and translation | 6.9E–05 | 131,133 |
| $J_{2A} = k_{2A} \times Q_1$ | Short-half-life protein synthesis | 2.6E–05 | 131,134,135,136-138 139 140 |
| $J_{2B} = k_{2B} \times Q_1 \times (1+Q_4)$ | Long-half-life protein synthesis | 1.6E–05 | 131,141 |
| $J_3 = k_3 \times Q_1 \times Q_{2B} \times Q_3 \times Q_5$ | Virus-RNA replication | 1.0E–01 | 33 |
| $J_4 = k_4 \times Q_3$ | Viral RNA translation | 6.9E–05 | 133 |
| $J_5 = k_5 \times Q_1 \times Q_{2B} \times Q_4$ | Recruitment of resources and host-cell protein machinery for virion assembly | 1.7E–03 | 33 |
| $J_6 = k_6 \times Q_1 \times Q_{2B} \times Q_4$ | Virion assembly | 3.0E–03 | 33 |
| $J_7 = k_7 \times Q_1 \times Q_{2B} \times Q_5$ | Virion budding | 8.3E–04 | 33 |
| $J_{13} = k_{13} \times Q_{2B} \times Q_1 \times Q_3 \times Q_5$ | Flow of host-cell resources diverted to let virus enter | 8.3E–02 | 33 |
| $J_{15} = k_{15} \times Q_1 \times Q_4 \times Q_{2B}$ | Flow of host-cell resources diverted to let virion assembly | 1.7E–03 | 33 |
| $J_{17} = k_{17} \times Q_1 \times Q_5 \times Q_{2B}$ | Flow of host-cell resources diverted to let virion shedding | 8.3E–04 | 33 |
| $J_{20A} = k_{20A} \times Q_{2A}$ | Flow of host-cell short-half-life proteins addressed to degradation | 3.9E–06 | 131, 142 |
| $J_{20B} = k_{20B} \times Q_{2B}$ | Flow of host-cell long-half-life proteins addressed to degradation | 1.6E–06 | 131,142 |
| $J_{21A} = k_{21A} \times Q_{2A}$ | Proteostasis mechanisms, including proteasome degradation and autophagy to re-cycle unfolded, old or not functional host-cell short-half-life proteins | 3.9E–06 | 131,142 |
| $J_{21B} = k_{21B} \times Q_{2B}$ | Proteostasis mechanisms, including proteasome degradation and autophagy of to re-cycle unfolded, old or not functional host-cell long-half-life proteins | 3.9E–06 | 131,142 |
| $J_{23} = k_{23} \times Q_1 \times Q_{2B} \times Q_3 \times Q_5$ | Flow of host-cell proteins recruited to let virus enter and RNA transcription | 8.3E–02 | 33 |
| $J_{25} = k_{25} \times Q_1 \times Q_{2B} \times Q_4$ | Flow of host-cell proteins recruited to let virion assembly | 1.7E–03 | 33 |
| $J_{27} = k_{27} \times Q_1 \times Q_{2B} \times Q_5$ | Flow of host-cell proteins recruited to let virion budding | 1.7E–03 | 33 |
| $J_{35} = k_{35} \times Q_4$ | Flow of viral RNA to embedd in the virion | 4.6E–05 | 33 |
| $J_{50} = k_{50} \times Q_1 \times Q_{2B} \times Q_5$ | Virion shedding | 4.6E–05 | 33 |

Equations representing the dynamics of the stocks and initial (calibration) values are expressed in ($x10^{12}$ ATP-eq) units.

**Supplementary Methods: the simulation code**

In our set of simulations, we modelled the temporal evolution of state variables (stocks) generated by the inflow and outflow of resources and quantified as embedded energy equivalents (ATP-eq). Scilab (https://www.scilab.org), an open-source scientific computation software, was used for such a purpose.

The architecture of our simulator is composed by three files, containing different scripts: CV_MainConsole, contains the main instruction for compiling the simulation, as well as the initial conditions. The file CV_system contains the set of differential equations, while CV_initiation contains the input equation, that describes the energy inflow to the system. The main file (CV_MainConsole) is contained in a folder, while the other two files are contained in a sub-folder, named ‘model’.

Data obtained from each simulation run, in form of files, with .cvs extension, are used to plot the results.

**Supplementary Methods Table 2**

**Code description and Script**

| File name | Script |
|---|---|
| ***CV_MainConsole*** | //Testing the COVID-19 model using literature data<br>clc<br><br>//Import CV functions<br>getd('model');<br><br>//Setting CV model parameter<br>//Stocks initial values refer to viral charge equivalent to 10k virions<br>//Parameters k refer to literature values<br><br>param = [];<br>param.k0 = 3.9e-6;<br>param.k1 = 6.9e-5;<br>param.k2A = 2.6e-5;<br>param.k2B = 1.6e-5;<br>param.k3 = 1e-1;<br>param.k4 = 6.9e-5;<br>param.k5 = 1.7e-3;<br>param.k6 = 3e-3;<br>param.k7 = 8.3e-4;<br>param.k13 = 8.3e-2;<br>param.k15 = 1.7e-3;<br>param.k17 = 8.3e-4;<br>param.k20A = 3.9e-6;<br>param.k20B = 1.6e-6;<br>param.k21A = 3.9e-6;<br>param.k21B = 3.9e-6;<br>param.k23 = 8.3e-2;<br>param.k25 = 1.7e-3;<br>param.k27 = 1.7e-3;<br>param.k35 = 4.6e-5;<br>param.k50 = 2e-3;<br>param.J = 0;<br>param.R = 1.7e2; |

| File name | Script |
|---|---|
| | param.E = 0;<br>param.H = 0;<br><br>//Setting initial conditions (TBD)<br>//Time start, end and steps resolution<br>Tbegin = 0;<br>Tend = 7;<br>Tstep = 1/86400<br><br>//Initial condition for stocks<br>Q1_0 = 3.9e0;<br>Q2A_0 = 1.3e1;<br>Q2B_0 = 1.1e1;<br>Q3_0 = 3;<br>Q4_0 = 2.4e-2;<br>Q5_0 = 0;<br><br>//Assigning ODE solver data<br>y0 = [Q1_0;Q2A_0;Q2B_0;Q3_0;Q4_0;Q5_0];<br>t0 = Tbegin;<br>t = Tbegin:Tstep:(Tend+100*%eps);<br>f = CV_system<br><br>//Solving the system<br>CV = ode(y0, t0, t, f); |
| ***CV_system*** | //set of ODEs for our COVID-19 model<br>function **CVdot**=CV_system(**t**, **CV**, **param**)<br><br>//fetching parameters<br>k0 = **param**.k0;<br>k1 = **param**.k1;<br>k2A = **param**.k2A;<br>k2B = **param**.k2B;<br>k3 = **param**.k3;<br>k4 = **param**.k4; |

| File name | Script |
|---|---|
| | k5 = param.k5;<br>k6 = param.k6;<br>k7 = param.k7;<br>k13 = param.k13;<br>k15 = param.k15;<br>k17 = param.k17;<br>k20A = param.k20A;<br>k20B = param.k20B;<br>k21A = param.k21A;<br>k21B = param.k21B;<br>k23 = param.k23;<br>k25 = param.k25;<br>k27 = param.k27;<br>k35 = param.k35;<br>k50 = param.k50;<br>R = param.R;<br><br>//Fetching solutions<br><br>Q1 = CV(1,:);<br>Q2A = CV(2,:);<br>Q2B = CV(3,:);<br>Q3 = CV(4,:);<br>Q4 = CV(5,:);<br>Q5 = CV(6,:);<br><br>//Evaluation of initiation<br><br>I = CV_initiation(param);<br>funcprot(0)<br><br>//Compute CVdot<br><br>//If replication is not enough, no outflow as virion<br><br>if Q3 < 3e-2 then k4 = 0 |

| File name | Script |
|---|---|
| | end<br><br>//If virions are not enough, no outflow as virion<br><br>if Q4 < 2.4e-4 then k5 = 0<br>end<br><br>//Computation<br><br>Q1dot = ((I)*(1+Q2A))+(k21A*Q2A+k21B*Q2B) - (k1*Q1) -<br>(k13*Q1*Q2B*Q3*Q5) - (k15*Q1*Q2B*Q4) - (k17*Q1*Q2B*Q5)<br>Q2Adot = (k2A*Q1) - (k21A*Q2A) - (k20A*Q2A)<br><br>Q2Bdot = (k2B*Q1*(1+Q4)) - (k21B*Q2B) - (k23*Q1*Q2B*Q3*Q5) -<br>(k25*Q1*Q2B*Q4) - (k27*Q1*Q2B*Q5) - (k20B*Q2B)<br><br>Q3dot = (k3*Q1*Q2B*Q3*Q5) - (k4*Q3)<br><br>Q4dot = (k4*Q3) - (k5*Q1*Q2B*Q4) - (k35*Q4)<br><br>Q5dot = (k6*Q1*Q2B*Q4) - (k7*Q1*Q2B*Q5)+(k35*Q4)<br><br>**CVdot** = [Q1dot; Q2Adot; Q2Bdot; Q3dot; Q4dot; Q5dot];<br><br>endfunction |
| ***CV_initiation*** | //Computes the initiation function for CV energetics<br>//Initiation function refers to the energy flow input to Q1<br><br>function **I**=CV_initiation(**Q1**, **Q2A**, **Q2B**, **Q3**, **Q4**, **Q5**, **param**)<br><br>//It returns the initiation for the CV model<br>//Fetching<br><br>k0 = **param**.k0<br>R = **param**.R |

| File name | Script |
| --- | --- |
| | //Define the function I<br><br>I = (k0*R); |
